# Visually-evoked choice behavior driven by distinct population computations with non-sensory neurons in visual cortical areas

**DOI:** 10.1101/2020.06.15.151811

**Authors:** Yuma Osako, Tomoya Ohnuki, Yuta Tanisumi, Kazuki Shiotani, Hiroyuki Manabe, Yoshio Sakurai, Junya Hirokawa

**Affiliations:** Laboratory of Neural Information, Graduate School of Brain Science, Doshisha University, 1-3 Tatara Miyakodani, Kyotanabe, Kyoto, 610-0394, Japan

**Author notes:** Contact Info: Correspondence (for Y.O.), (for J.H.).

## Abstract

It is widely assumed that variability in visual detection performance is attributed to the fidelity of the visual responses in visual cortical areas, which could be modulated by fluctuations of internal states such as vigilance and behavioral history. However, it is not clear what neural ensembles represent such different internal states. Here, we utilized a visual detection task, which distinguishes perceptual states to identical stimuli, while recording neurons simultaneously from the primary visual cortex (V1) and the posterior parietal cortex (PPC). We found distinct population dynamics segregating hit responses from misses despite no clear differences in visual responses. The population-level computation was significantly contributed by heterogenous non-sensory neurons in V1, whereas the contribution from non-neurons with the previous outcome selectivity was prominent in PPC. These results indicate different contributions of non-sensory neurons in V1 and PPC for the population-level computation that enables behavioral responses from visual information.

## Introduction

Identical sensory stimuli sometimes evoke different perceptual and behavioral responses. For instance, in a sensory detection task, human or animal subjects are instructed or well-trained to reliably report the presence and absence of sensory stimuli to obtain rewards. When the sensory evidence is near the threshold for the decision criterion, the subjects’ reports vary across trials despite subjects’ best efforts to get rewards. Interestingly, even if they report the absence of stimuli, it is sometimes possible that they could correctly guess the contents of the stimuli above chance level if they are forced to answer^1–4^. Revealing the neural mechanisms underlying such trial-by-trial variability of perceptual reports is crucial to understand how the brain exploits sensory information for optimal decision making.

The trial-by-trial variance of the responses to identical stimuli is believed to reflect noises in sensory information conversion into motor outputs^5^. It has been demonstrated that the variability of firing rates of sensory neurons is responsible for the trial-by-trial variability of choices^6,7^. However, the accumulating evidence suggests that perceptual decision is also significantly affected by latent subjective states reflecting task engagement^8^. For instance, behavioral response variability is correlated with mind wondering in humans^9^ and fluctuations of physiological and behavioral states in animals^10–13^. These drifts of subjective states could be partially attributed to the fluctuation of cortical states^13–15^. The synchronization and desynchronization of many neurons in particular areas of the cortex could affect the efficiency of the population coding^16,17^. Related to this, shared response variability in pairs of sensory neurons (i.e., noise correlation), modulated by attention, arousal, and reward expectation, can affect efficient coding of stimulus features and sensory processing, resulting in behavioral variability in a sensory detection task. Besides, the task engagement is known to be modulated by the trial-by-trial experience of decisions and its outcomes^18–24^, which is supported by distinct populations of neurons in association areas^25–27^. Furthermore, some studies suggest that neurons that do not explicitly respond to a stimulus contribute to the texture discriminations in the somatosensory cortex^28^, working memory coding in the prefrontal cortex^29^, stimulus/choice coding in the auditory cortex^30^ and category representation in the prefrontal cortex^30^. These studies highlight the potential contribution of non-sensory neurons in modulating sensory processing resulting in trial-by-trial variability of responses. However, it has been elusive how non-sensory neurons are coordinated with sensory neurons for optimal sensory decisions.

Many previous studies revealed that neural activity in the primary visual cortex (V1) and posterior parietal cortex (PPC) plays a crucial role in visual detection behavior^31,32^. Patients with V1 lesion report subjective blindness^33,34^ and direct optogenetic inhibition of rodent V1 impaired visual detection behavior^35^. On the other hand, PPC is known to play essential roles in selective attention and reward-history bias^27,36^ and regulates response properties of V1 neurons^37–39^. Recent imaging studies examined visual perceptual behavior during the go/no-go detection task and found that task requirements^40^ heavily modulates visual responses in PPC, and heterogenous recruitments of V1 neurons play an important role in visual detection^40,41^. Together, these studies support the notion that V1 and PPC form distinct cortical states at a population level that integrates task-relevant external signals^42^ with internal states for subjective detection performance.

Though neural imaging studies addressed population coding of sensory processing across different cortical areas, the go/no-go task paradigm, which is often used in those experiments with head-fixed animals, is susceptible to subjective biases: go trials may contain false alarms, and no-go trials may contain false rejections^43^ due to fluctuating internal states as described above. To further classify such internal states during the visual detection task, we previously developed a spatial-visual cue detection task for free-moving rats^44^. The task combines a two-alternative spatial choice task with a third option for void stimulus, which allowed us to differentiate three distinct visual detection states based on choice types during the task. By taking advantage of these relatively homogenous trials with different choice types to identical stimuli, we revealed distinct contributions of V1 and PPC neurons at the population level for those choice types. Distinct coordinated activities differentiate these choice types from non-sensory neurons, whereas the fidelity of the visually-evoked activities in V1 and PPC did not. In particular, non-sensory neuronal subpopulations sensitive to the previous outcome in PPC relatively contributed to the population coding compared to sensory neurons. On the other hand, the recruitment of V1 neurons were more distributed. Besides, response variability between neurons (i.e. noise correlation) segregated choice types in V1, which was partially influenced by previous outcome. Our findings provide the notion that non-sensory neurons modulate the accurate visual decision in a history-dependent manner.

## Results

### Rats performed visual detection task based on their internal threshold

We trained seven rats to perform a spatial visual-cue detection task (Fig.1a), which is essentially a 3-alternative choice design and encourages animals to report the presence and the absence of the peripheral visual stimuli as described in our previous study^44^. Briefly, the rat initiated a trial by nose-poking to the central port. They were rewarded by choosing peripheral ports when a visual stimulus is presented (left or right) or keeping the nose in the central port when no peripheral stimulus is presented. In half of the trials, the central port is closed 0.5 s after stimulus presentation timing to force animals to make peripheral choices.

**Figure 1.**
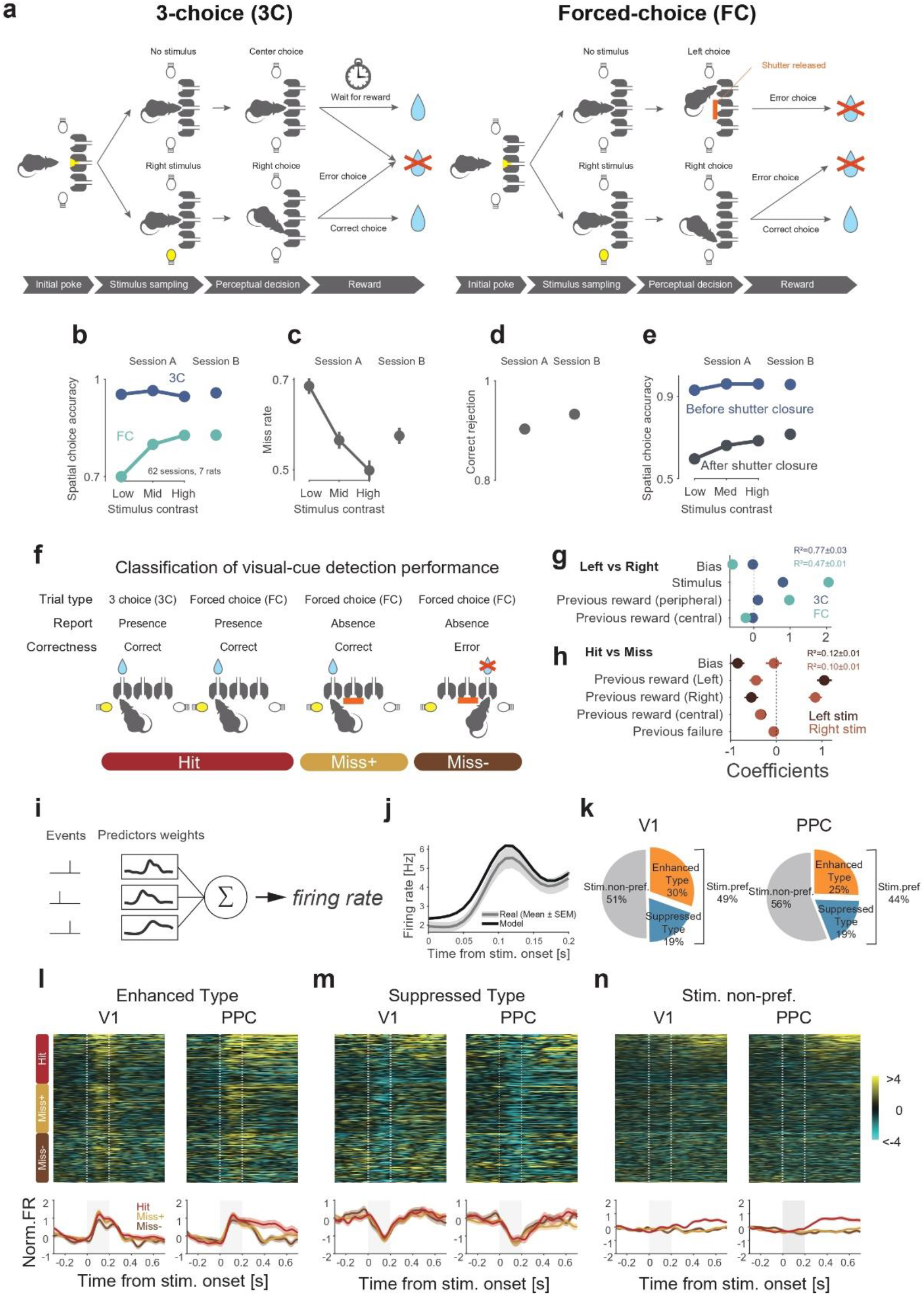
Neural recording in V1 and PPC during spatial visual-cue detection task. **a,** Schematics of behavioral paradigm. Rats initiated a trial by nose-poking into the central port and waited for 0.2-0.6s to receive a peripheral stimulus. Rats were rewarded by poking to the corresponding peripheral port when the peripheral stimulus was presented. Rats were rewarded in the central port when the peripheral stimulus was not presented. FC and 3C trials were identical except that the central port was shut in FC trials when rats kept nose-poking in the central port more than 0.5s. **b**, Spatial choice accuracy in 3C and FC trials with graded visual contrast in session A and a fixed contrast in session B. **c**, Miss rate in 3C trials. **d**, Correct rejection rate in 3C trials. **e**, Spatial choice accuracy before and after shutter closure in FC trials. **f**, Conceptual illustration of three distinct choice types. We classified spontaneous correct response, forced correct response and forced error choice as Hit, Miss+ and Miss-, respectively. **g-h**, The impact of task parameters on behavioral variability using GLM fitting with several task parameters to Left/Right choice in 3C and FC (**g**), and to Hit/Miss choice in peripheral stimulus trials (**h**). **i**, Illustration of kernel regression approach to fit the neural activity by three task predictors. **j**, Example trial-averaged responses (plot with the shaded area) and model predictors (Thick line). **k**, Fraction of stimulus preferring and non-preferring neurons in V1 (left) and PPC (right). **l-n**, Trial-averaged neural activity for each choice type in Enhanced type stimulus preferring neurons, Suppressed type stimulus preferring neurons, and stimulus non-preferring neurons in V1 and PPC. All responses were z-scored, and neurons were sorted by max peak latency in Hit trials.

To confirm whether rats have a steady internal detection criterion, we alternated the prove sessions with graded stimulus strength (session A) and the neural recording sessions with a constant near-threshold stimulus strength (session B). In session A, as consistent with the previous study, the peripheral choice accuracy in forced-choice (FC) trials decreased from approximately 80% to 65% as the visual contrast decreased (Fig.1b, blue, Extended Data Fig.1), while the accuracy was maintained at >90% regardless of visual contrast in the three-choice (3C) trials (Fig. 1b, sky blue, Extended Data Fig.1). The rats missed the visual stimuli more often (from 50% to 70%) as the visual contrast decreased (Fig.1c, Extended Data Fig. 1). They also showed >90% correct rejection performance when the visual stimulus was not emitted in 3C trials (Fig.1d, Extended Data Fig. 1). These results confirmed that rats have a generalized strategy to make peripheral choices across stimuli only when their internal detection criterion is met. Also, rats showed correct peripheral choices above the chance level when the shutter forced them to make peripheral choices after choosing to stay in the central port (Fig.1e, gray, Extended Data Fig.2). Thus, rats received visual information but did not exploit it maximumly. We labeled trials with different choice types as Hit, Miss+, and Miss-according to the choice performances (Fig.1f). To find out the influences of trial history in those choice types, we conducted a generalized linear model (GLM) analysis for spatial choice (left and right) and hit/miss choice (Fig.1g-h). In FC trials, about half of the choice variance was explained by the model showing significant influences from previous peripheral rewards and the current stimulus direction. In contrast, in 3C trials, influences of rewards disappeared, and 77% of the choice variance was accounted by stimulus alone (Fig.1g). In addition, a relatively small portion of the total variance (10%) for hit/miss choices was explained with the model containing past rewards, in which the major contribution was previous reward ipsilateral to the current stimulus (Fig.1h). Thus, our task segregated trial-by-trial variability of responses during the detection task into distinct categories (Hit, Miss+, Miss-).

### Stimulus-preferring neurons in V1 and PPC were activated regardless of choice types

We recorded neurons simultaneously from the right V1 (N_V1_ = 515 neurons) and PPC (N_PPC_ = 436 neurons) using chronic tetrode implants during the task performance (Extended Data Fig.3). To investigate how visual neuronal responses contribute to the behavioral responses, we first identified stimulus preferring neurons using time-locked kernel regression analysis with multiple task predictors such as contra-, ipsi-stimulus, pre-movement, and pre-choice kernels (Fig. 1i-j, Extended Data Fig.4a-c, see Methods). Half of the neurons were defined as selective to visual stimulus in V1 (49%, N = 256) and PPC (44%, N = 192) (Fig. 1k, Extended Data Fig.4e-f). Based on the results, we classified neurons as enhanced type sensory neurons, suppressed-type sensory neurons, and the rest as non-sensory neurons.

Both V1 and PPC neurons showed stimulus-dependent activity (enhanced and suppressed from pre-stimulus baseline) regardless of the choice types (Fig.1l-n), even if rats did not make peripheral choices (Miss+ and Miss− trials). Classification analysis of stimulus (presence/absence) showed that stimulus preferring population decoded stimulus presence near-perfectly regardless of choice types, whereas stimulus non-preferring population did not predict it (Extended Data Fig.4f). The results confirmed that the stimulus period’s activity is evoked by visual stimulus but not by stimulus expectations. These results indicate that missed responses are not sufficiently explained by the absence of visual information in the visual cortex. Instead, all the choice types showed a similar time course of mean firing rates in stimulus-preferring neurons, particularly in V1, at least, until the initial peak (around 100 ms from stimulus onset) of the stimulus responses (Fig. 1l-m, Extended Data Fig.4g). The time course of stimulus non-preferring neurons in both V1 and PPC also did not show apparent differences on average among choice types until 0.2s after stimulus presentation where behavioral responses occur (Fig.1l-m).

### Significant contributions of non-sensory neurons in V1 as well as PPC for separating different choice types as population activity

Next, we address what neural population states differentiated self-initiated correct responses from withholding correct responses. To this end, principal component analysis (PCA) was applied to the pre- and post-stimulus presentation window (−0.1 – 0.15s after stimulus onset) calculating each neuron’s weights, which is optimized to capture variance of neural activity across the choice types and time. We identified three dimensions that captured 73, 73% for the whole population in V1 and PPC, respectively (Extended Data Fig.5c-d). As a result, population dynamics showed unique trajectories for Hit and Miss+ (Fig.2a). To estimate the difference of neural trajectories, we compared PC projections between Hit and Miss+ in each PC and used a sensitivity index (d’) between Hit and Miss+ calculated for the first three PCs (see Methods). We found that significant separation between choice types at the analysis window in both V1 and PPC (Fig.2d), suggesting Hit and Miss responses are the results of the coordinated activities of many neurons.

**Figure 2.**
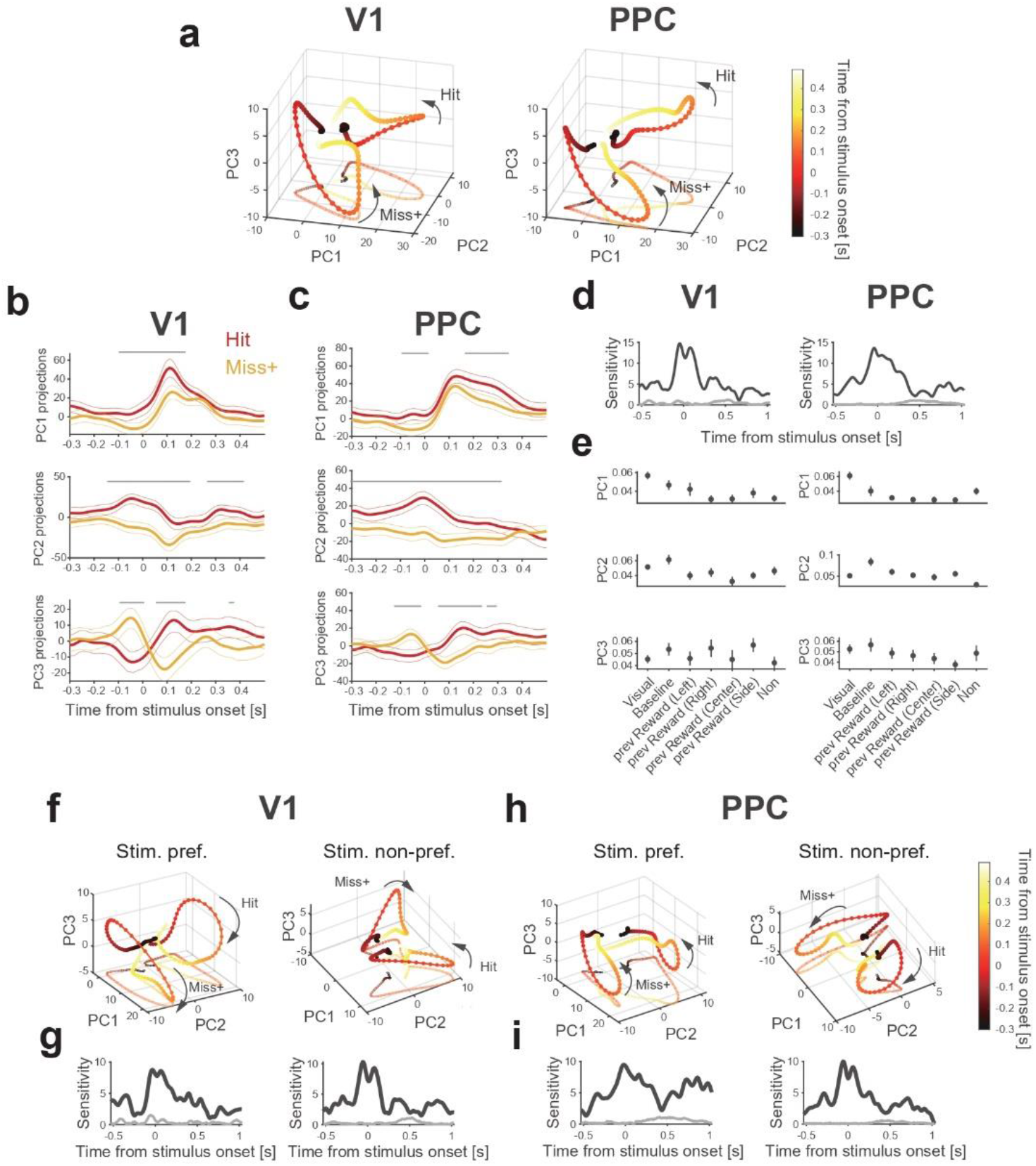
Population dynamics in pre-stimulus and stimulus subspaces in V1 and PPC. **a,** Population responses of the whole population projected onto three dimensions of the analysis window (−0.1-0.15s from stimulus onset) in V1 and PPC. Each color corresponds to time relative to stimulus onset. **b**, **c**, PC projections of **a**, **b** in each principal component. Each plot was shown as Mean ± 95% confidence interval. Gray lines above each projection are significantly different across choice types. **d**, Sensitivity index (d’) across choice types in different time window for V1 and PPC. Black line indicates real data and thin line indicates shuffled data. **e**, Absolute value of weights in each PCs were shown as Mean ± SEM. **f**, **h**, Population dynamics of stimulus preferring and non-preferring population projected onto three dimensions of the analysis window (−0.1-0.15s from stimulus onset) in V1 and PPC. **g**, **i**, Same as in **d**, but calculated from stimulus preferring and non-preferring population.

Interestingly, V1 and PPC population activities contain similar principal components that reflect different task variables. For instance, PC1 shows distinct dynamics peaked around visual stimuli onset (Fig.2b), of which amplitude separates the choice types. The component has a major contribution from visual-responsive neurons (Fig. 2e). On the other hand, PC2 separates Hit and Miss+ trials robustly long before stimulus onset (Fig.2b), and that is mainly supported by neurons with distinct baseline activities (Fig. 2e). Finally, PC3 shows theta-range oscillatory components, of which phase separates choice types, suggesting a significant influence of global fluctuation of activity across V1 and PPC on Hit/Miss+ behavior. These results indicate that Hit and Miss+ behaviors are mediated by multiple levels of processing from a variety of neurons at pre- as well as during-stimulus epochs.

To confirm this activity pattern resides in different population types, we repeated the same analysis dividing the population into stimulus preferring and non-preferring populations (capturing 78, 78% variance in stimulus preferring population and 70, 68% variance in stimulus non-preferring population in V1 and PPC, respectively). In both V1 and PPC, separative activity between choice types was observed in both stimuli preferring and non-preferring populations (Fig.2f-i). We also found the three types of dynamics in stimulus non-preferring neurons (ExtFig.5a,b). In particular, theta-range oscillatory dynamics, of which phase differentiate choice types, was only evident in the non-preferring population. These results emphasize the important role of stimulus non-preferring neurons as well as stimulus preferring neurons, which consists of distinct temporal components largely in common between V1 and PPC.

### Population representation of choice types is distributed across heterogeneous individual neurons

So far, we have addressed distinct population activities across choice types in pre- and during-stimulus windows. Given this finding, we asked whether the neural computation contributing to the choice types is only composed of individual neurons that contain different choice type information, or neurons that have subtle information. We first used a linear support vector machine (SVM) algorithm to predict the choice types (Hit/Miss+) of trials from the single neuron activities in the stimulus epoch (0 – 0.2s from stimulus onset) (Fig.3a, upper). Only a few neurons had a better classification accuracy than shuffled classification accuracy in both V1 and PPC (Fig.3a). This indicates that most of the single neurons do not have sufficient information to separate choice types. We next addressed whether the only subset of neurons with high selectivity to the choice types contributed to computation at the population level or whether the other neurons with low selectivity also contributed to population computation. We sorted the neurons based on the individual decoding accuracy in ascending order, and gradually incorporated all individual neurons. The classification accuracy improved in both V1 and PPC by incorporating neurons with low information of choice types (Fig.3b, Thick line: subpopulation decoding accuracy, Thin line: Individual max decoding accuracy). These results indicate that information of individual neurons for separating choice types is poor, but including these single neurons gradually improved the classification of choice types. The same analysis was conducted for the pre-stimulus epoch (−0.2 – 0s from stimulus onset), indicating classification accuracy was improved with the incorporation of neurons (Fig.3c-d).

**Figure 3.**
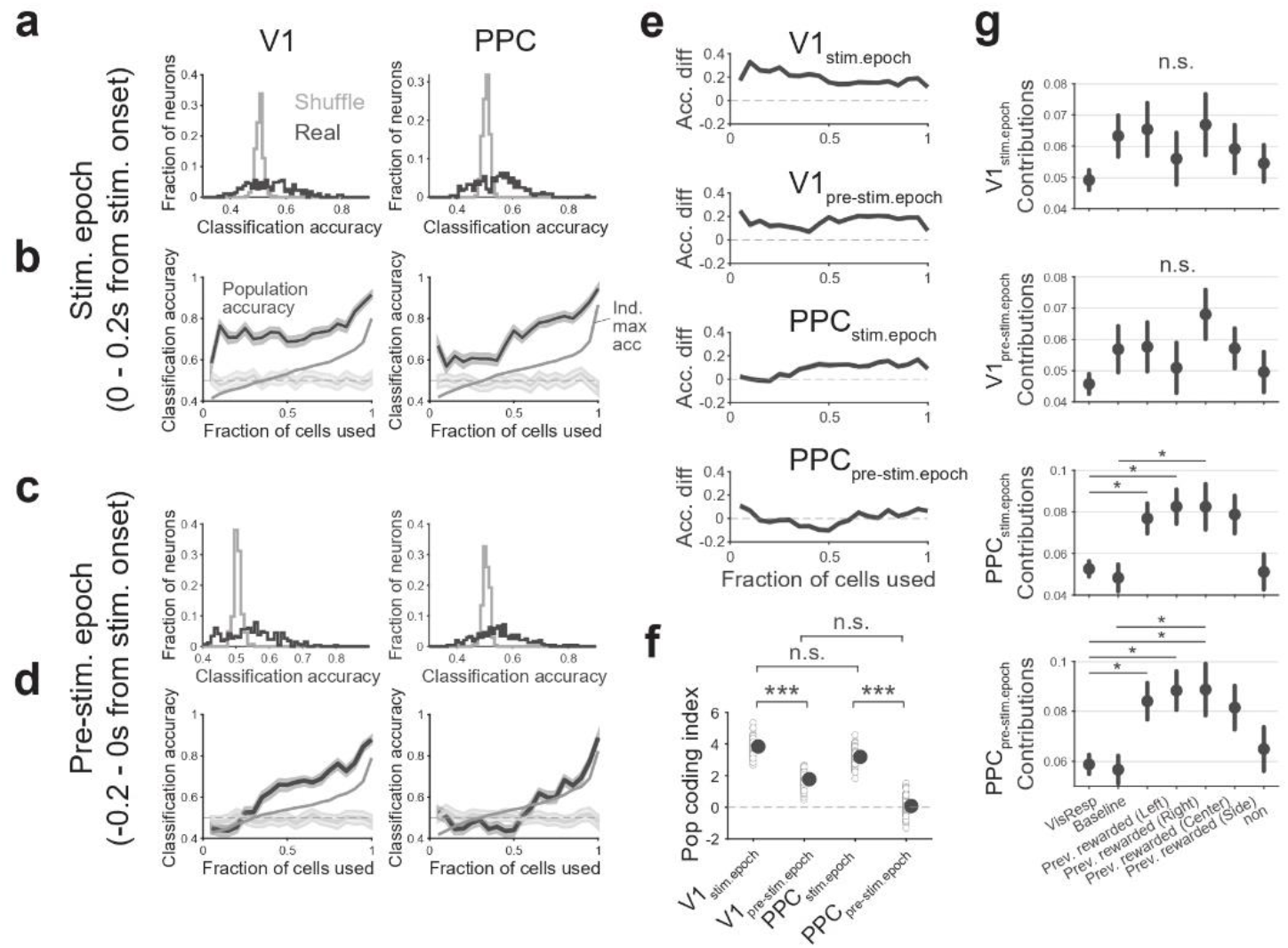
Distributed representation by sequential population size. **a**, Histogram of classification accuracy for choice types for individual neurons (thick line) and shuffled data (thin line) based on the normalized firing rate in stimulus epoch (0-0.2s from stimulus onset) in V1 and PPC. **b**, Classification accuracy for choice types by incorporated population sizes. Neurons we included by descending order based on the histogram from the whole population. Each line was shown as Mean ± 95% confidence interval. **c**, **d**, Same as in **a**, **b**, but in pre-stimulus epoch (−0.2-0s from stimulus onset). **e**, Difference of classification between population accuracy and individual max accuracy. **f**, Population coding index for each epoch in V1 and PPC. **g**, Absolute value of weights for different neurons types in classifier trained on all neurons. Each line was shown as Mean ± SEM. *p < 0.05, one-way ANOVA followed by post-hoc Tukey tests. **h**, Classification accuracy of classifiers for each pair of training/testing time points for the whole population in V1 and PPC. **i**, Same as in **h**, but for stimulus preferring and non-preferring

To quanti fy the degree of improved classification accuracy by the cnordinated activity of multiple neurons relative to that by the individual neurons, we compared the classification accuracy between a subpopulation and the neuron with maximum classification accuracy within the subpopulation (Fig.3e-f). Both V1 and PPC showed steady improvement of the classification accuracy by the population effect in the stimulus epoch, whereas, during the pre-stimulus epoch, there were no apparent improvements of the classification accuracy in V1 and PPC, suggesting population-level computations driven by stimulus presentation.

Next, we addressed how neuron types (i.e., stimulus preferring and non-preferring) contribute to these population-level computations in pre-stimulus and stimulus epoch. We calculated classifier weight magnitudes of stimulus preferring and non-preferring neurons (Fig.3g). While there is no difference in the weight magnitudes in pre-stimulus and stimulus epochs in V1, weight magnitudes for the sensory non-preferring neurons, especially with the contributions from the previous-outcome preferring neurons, were significantly higher than those for the sensory preferring neurons in PPC. These results indicate that sensory preferring and non-preferring neurons in V1 are similarly informative for choice types, while the non-preferring population in PPC, especially neurons selective to previous outcomes, play a significant role in differentiating choice types.

So far, we have shown that the sensory preferring and non-preferring neurons contribute to the population-level computation with distinct manners between V1 and PPC (Fig.3g). Are those subpopulations temporally coupled to each other as a whole-population computation? If they are, neural representation for each subpopulation will follow a similar pattern over time. To address this possibility, we trained the classifiers for each time epochs, and then applied it to all-time epochs to calculate the temporal classification performance for whole and subpopulations (Fig.4a-b). As consistent with previous studies^45^, the classifiers from the whole population were stable across time in both areas, especially in PPC (Fig.4a-b). High classification accuracy 0.2 sec after stimulus onset simply reflects the different actions between Hit and Miss+, showing distinct decodability from other time points. The classification accuracy patterns were qualitatively different between stimulus-preferring and non-preferring populations in both V1 and PPC, suggesting uncoupled computation between those subpopulations. Interestingly, there was no sign of increased classification accuracy at the timing of visual responses around 0-0.2 sec in stimulus-preferring populations, suggesting that different choice types are not triggered by visual neuronal responses in V1 nor PPC, but rather by the preparatory state of the neural populations. To estimate the stability of contributions of neurons for the decoding of choice types across times, we calculated Pearson’s correlation coefficients of neural weights of pairs of classifiers at different times (Fig.4c-d). If population computation is similar across different times, correlation coefficients will be tolerant of decaying. For a comparison between whole populations in V1 and PPC (Fig.4f top), PPC was relatively stable compared to V1. Such a stable population computation was, in particular, evident in the non-preferring population in PPC (Fig.4f below), whereas the stability was lower in both stimulus-preferring and non-preferring populations in V1. Together, these results suggest that sensory preferring and non-preferring neurons are not likely to coordinate each other for the population-level computations, and the sensory non-preferring neurons in PPC provide a significant contribution to the long decay constant for the population coding.

**Figure 4.**
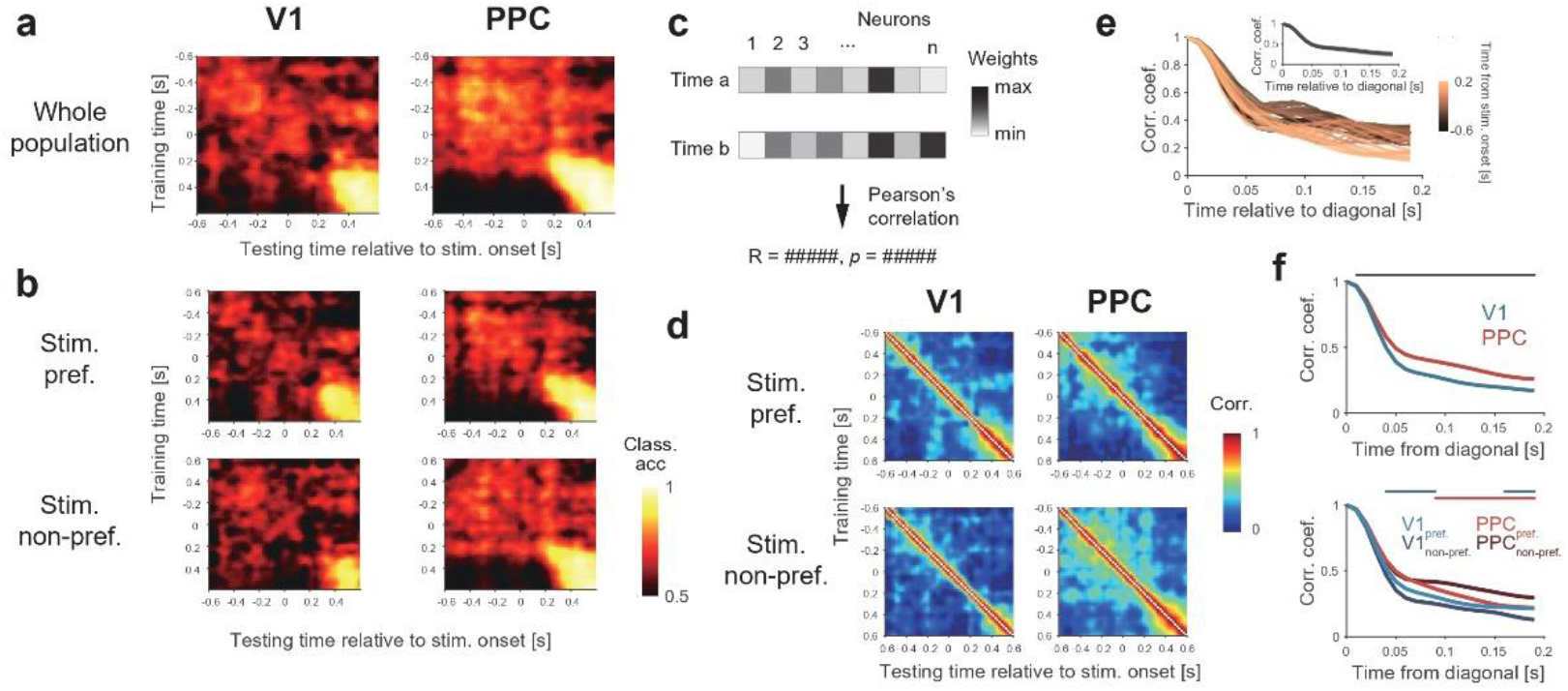
Time-varying classification performance and weight correlation. **a**, Classification accuracy of classifiers for each pair of training/testing time points for the whole population in V1 and PPC. **b**, Same as in **a**, but for stimulus preferring and non-preferring population. **c**, Schematics of calculation for weight correlation. **d**, Weight correlation of classifiers for each pair of training/testing time points for sub-population in V1 and PPC. **e**, Time-varying correlation coefficient of classifiers in each time (−0.6 – 0.2s from stimulus onset). Inset shows mean time-varying correlation coefficients. The shaded area indicates the standard deviation (S.D.). **f**, Time-varying correlation coefficients in each population. Lines on top of panels indicate significant differences between V1 and PPC (Black), stimulus preferring and non-preferring in V1 (Cyan) and in PPC (Red). p<0.05, Wilcoxon rank sum test for V1 and PPC, oneway ANOVA followed by post-hoc Tukey tests for subpopulations.

### V1 Noise correlation was increased in forced detection performance before and after stimulus presentation

Our results thus far demonstrate that both stimulus preferring and non-preferring neurons are essential for choice types in V1 and PPC. However, we used “pseudo-population” that combined neural activity recorded in different trials in these analyses. Therefore, our analysis missed considering the correlation structure of pairs of simultaneously recorded neurons within each trial (i.e., noise correlation). If the noise is closer to random across neurons (low noise correlation), information coding is more reliable and efficient^46^. We first examined classification accuracy in each session using a simultaneously recorded population. Only the V1 population showed significantly higher classification than shuffled data (Fig.5a, p = 0.0460, p = 0.1985 in V1 and PPC, respectively). Next, we compared classification accuracy with a decorrelated population in which each neuron in the same session was randomly taken from different trials. Only the V1 population was significantly decreased classification accuracy in the decorrelated population (Fig.5b), indicating that the correlation structure, at least, in V1 was crucial for population computation.

**Figure 5.**
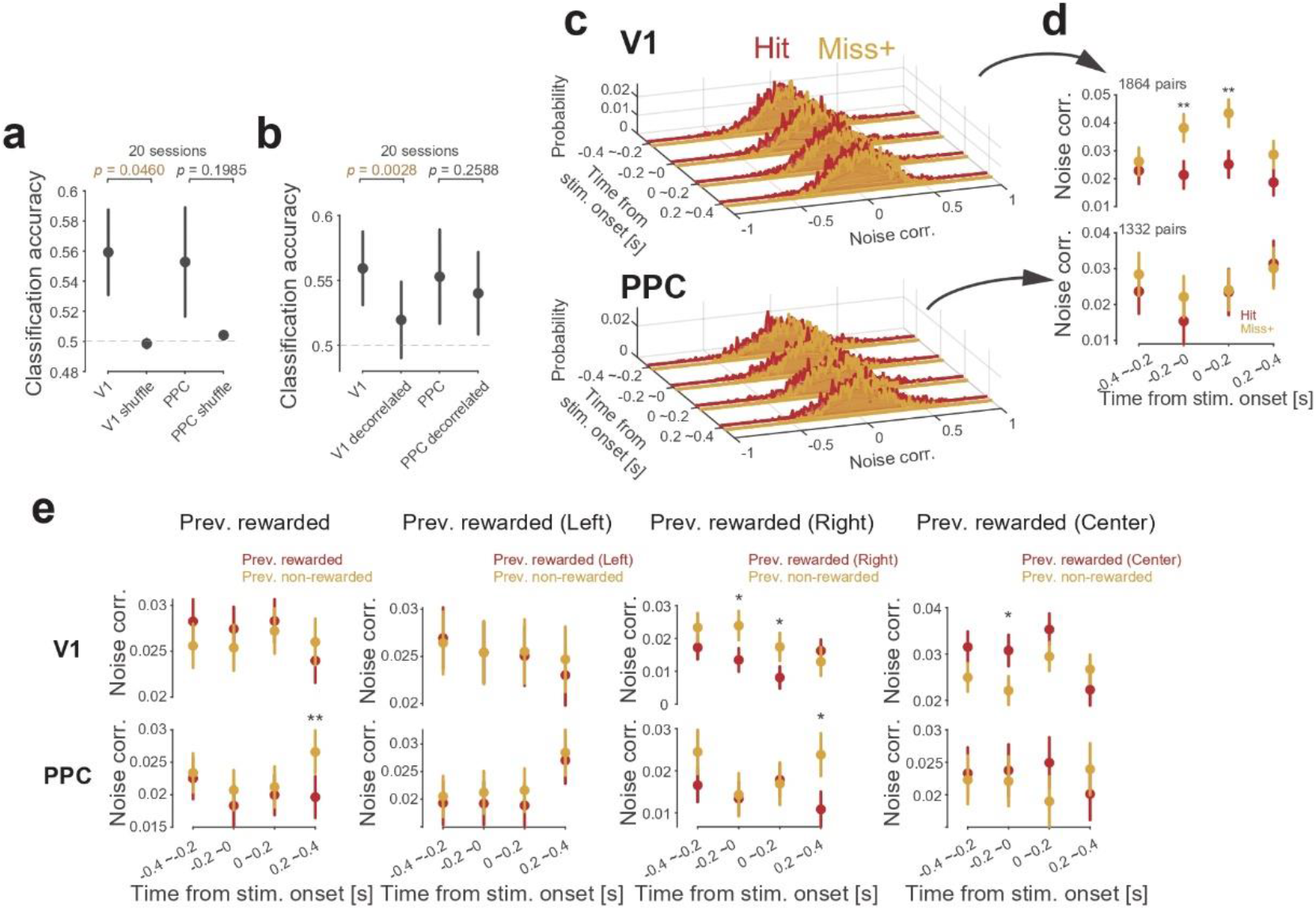
Noise correlation in Miss+ trials was increased around stimulus presentation in V1. **a**, Classification accuracy for the real and shuffled population for each session in V1 and PPC. p < 0.05 in V1, paired t-test. **b**, Classification accuracy for the real and de-correlated population for each session in V1 and PPC. p < 0.05 in V1, paired t-test. **c**, Histograms of noise correlation of simultaneously recorded neuron pair in Hit and Miss+ in V1 and PPC. **d**, Mean noise correlation in each time epoch in V1 and PPC. **p<0.01, t-test. **e**, Noise correlations in previous rewarded and non-rewarded trials for each 200 ms time epochs relative to stimulus onset in V1 and PPC were shown as mean ± SEM. *p < 0.05, **p < 0.01, paired t-test.

To investigate the correlation structure in V1 and PPC for choice types, we calculated noise correlations in 200ms sliding time windows relative to stimulus onset (Fig.5c, −0.4 – 0.4s from stimulus onset). We found that noise correlation in Miss+ trials increased in −0.2s – 0.2s from stimulus onset compared to Hit in V1 neuron pairs, while PPC neuron pairs did not differ in choice types (Fig.5d). Such a difference in the noise correlation was only apparent in neuron pairs between regular-spiking neurons (Extended Data Fig.6d). To examine whether reduced noise correlation is associated with previous outcomes, we compared the noise correlation between previous rewarded and unrewarded trials. We found that noise correlation in V1 was significantly reduced in previous ipsilateral reward trials around stimulus presentation timing (Fig.5e), while that was significantly increased when animals were previously rewarded in the central port. Such a reduction of the noise correlation after rewarded trials were not observed in PPC excepted for movement epoch (0.2 - 0.4s from stimulus onset). These results suggest that neuronal coupling in V1 plays an important role in accurate visual detection performance and the reward history partially responsible for modulating coordination of neurons for subsequent trials.

## Discussion

It is widely believed that the trial-by-trial variance of visual detection performance is attributed to the fidelity of visual responses in sensory neurons. Our results demonstrated significant contributions of the non-sensory neurons in V1 and PPC for reliable visual detection performance. The near-threshold stimuli used in our task-induced trial-by-trial variability in visual detection performance, and we further classified them into three types of the trial with distinct perceptual states, which are not differentiated in the previous studies. Surprisingly, both Hit and Miss+ trials show comparable levels of visual responses at the single neuron levels. Instead, we showed multiple evidence for population-level computation that contributes to the correct responses to visual stimuli (Hit/Miss+) in V1 and PPC. First, despite no apparent differences in averaged activity among choice types (Fig.1l-n), we found a specific divergence between choice types at the multiple levels of the population activity, particularly with the robust contribution from the stimulus non-preferring neurons in both V1 and PPC (Fig.2). Second, during pre-stimulus and stimulus epochs, contributions of individual sensory non-preferring neurons for classification of choice types were significantly greater compared to the stimulus preferring neurons in PPC but not in V1 (Fig.3g). Third, we found that individual neurons that poorly represented choice types also contributed population coding of choice types especially in stimulus presentation timing (Fig.3a-f). Fourth, these contributions were more stable in stimulus non-preferring population in PPC but not in V1 (Fig.4). Finally, V1 neuron pairs, but not PPC, showed increased noise correlation in Miss+ trials before and during visual stimulus presentation (Fig.5c-d), indicating that the V1 population gains efficiency of information processing for future behavioral output.

It has been postulated that stochastic behavioral responses to identical sensory stimuli are generated by fluctuations of background neural ensembles preceding to the external inputs^47^. Our model-free analysis revealed that there are at least three distinct population dynamics that differentiate behavior to identical stimuli (Fig.2), all of which have significant contributions from non-sensory neurons. First, there were transient dynamics of non-sensory neurons after stimulus presentation in segregating choice types in both V1 and PPC. Such population activity by non-sensory neurons was not coupled with stimulus-preferring subpopulation activities (Fig.4b), suggesting non-sensory neurons are not directly modulating sensory responses in V1. Second, there were slow dynamics of baseline activities that significantly contributed to separating the choice types. Finally, we identified theta-range oscillatory components, which were observed, especially in non-sensory neurons of both V1 and PPC, as significant factors that differentiate choice types. Such populational fluctuation could play an important role in coordinating many neurons to generate appropriate behaviors^47^. The results agree with the hypothesis that correlated noise in sensory areas, partially driven by selective spatial attention from previous outcomes, filter sensory inputs from a noisy environment^46,48^. Our data demonstrated distinct population dynamics that would be generated by different neural mechanisms.

Relatively weak contribution of the visually-evoked activities in V1 population coding could be a unique feature of our task design because the animals have a third option, of which experienced reward value may suppress peripheral choices even when the animals recognize the stimuli. However, it should be noted that ignoring the presence of stimuli is never rewarded in our task and, thus, clearly suboptimal behavior. In addition, our data show that the rewards in the previous trials only partially explain the behavioral variance in Hit/Miss choices (Fig.1h) as well as the variance in noise-correlation (Fig.5e) in V1 and population coding in PPC (Fig.3g). Therefore, we conclude that the value-based decisions do not solely explain the recruitments of non-sensory neurons in V1 and PPC. On the other hand, it is possible that irreverent movements during the task performance could have affected Hit responses due to suboptimal head/body positions^49,50^. However, our data shows the noise-correlation in V1 is specifically increased in Miss+ trials, suggesting that, at least, the accurate performance in Hit trials is due to an intrinsic population-level mechanism that can be related to sensory-motor transformation during the task^40^. Although it has been indicated the significant contributions of non-sensory neurons for perceptual decision makings^28–30^, most of them employ sensory categorization tasks in the forced-choice paradigm, which unavoidably suffers from stimulus uncertainty causing subjective biases due to value-based decisions. Our data support and extend these findings by showing that even in the simplest sensory detection task, which does not have inherent stimulus uncertainty and is less contaminated by value-based decisions, non-sensory neurons in V1 and PPC plays a significant role in sensory decisions at a population level.

## Acknowledgments

We would like to thank members of the Laboratory of Neural Information at Doshisha University for helpful discussion. This research was supported by the JSPS KAKENHI Grant Numbers 19J21544 (to Y.O.), 20H00109, 20H05020 (to Y.S.), 19J12634 (to T.O) and 16K18380, 19H05028, 19K06948 (to J.H.).

## Author Contributions

Y.O., T.O., Y.S., and J.H. designed the experiments. Y.O. performed the experiments and mainly analyzed the data. T.O., Y.T. and K.S. analyzed the data. H.M, Y.S. and J.H. supervised the project. All authors contributed to writing the manuscript.

## Methods

### Subjects

Seven male Long-Evans rats (Shimizu Laboratory Supplies, Kyoto, Japan) weighing 200-268 g at the beginning of the training were individually housed and maintained on a laboratory light/dark cycle (lights on 8:00 A.M. to 9:00 P.M.). Rats were placed on water restriction with *ad libitum* access to food. The animals were maintained at 80% of their baseline weight throughout the experiments. All experiments were implemented in accordance with the guidelines for the care and use of laboratory animals provided by the Animal Research Committee of the Doshisha University.

### Behavioral apparatus

The behavioral apparatus (Fig.1a) has been previously described^2^. An operant chamber (O’Hara, Tokyo, Japan) with three ports in the front wall for nose-poke responses was enclosed in a soundproof box (Brain Science Idea, Osaka, Japan). Each port was equipped with an infrared sensor to detect the animals’ nose-poke responses. Visual cues were presented using white light-emitting diodes (LEDs) (4000 mcd; RS Components, Yokohama, Japan) placed on the left and right walls of the operant chamber, as in Fig.1a. Water rewards were delivered from gravity-fed reservoirs regulated by solenoid valves (The Lee Company, Westbrook, CT) through stainless tubes placed inside of the left and right target-ports. Stimulus and reward deliveries were controlled with Pulse Pal^41^, and behavioral responses were measured with Bpod (Sanworks, Stony Brook, NY).

### Visual cue detection task

The visual cue detection task design was described previously^2^. The task was comprised of randomly interleaved three choice (3C) and forced-choice (FC) trials with equal probabilities in a session. The only difference between the trial types was that, in FC trials, the central port was shut with the shutter door to prevent the rat from continuing to central nose poke (Fig. 1a). After a fixed 2.5 s inter-trial interval (ITI), the central port was illuminated by an interior LED of the central port signaling the ready state of trial initiation. The rats initiated each trial by making nose pokes into the central port. After a 0.2–0.6 s random stimulus delay, the visual stimulus was presented from the left or right side for a duration of 0.2 s. Rats were allowed to make a choice response after the end of the stimulus delay period. The trials where animals prematurely left the port before stimulus delay were canceled and they needed to re-initiate the trials. We randomly provided one of three levels of visual brightness (difficult, medium, and easy) for each trial by modulating the voltage ranging 0.02–5.1 lx in session A. Difficult and easy stimuli were selected for all subject based on the minimum and maximum LED voltage dynamic range. Medium stimuli were chosen for each subject such that the subject detected stimuli with medium accuracy between easy and difficult stimuli in the forced-choice (FC) trials. The probabilities for left, right, or no visual stimulus were equal (33% per condition) in 3C and FC trials. The reward was given if rats chose the same side where the visual stimuli were emitted in the 3C and FC trials. If animals kept nose poke more than X s in the central port after the presentation of the visual stimuli, the trial was treated as a miss error. X was drawn from the uniform distribution with a range of [0.5, 1]. Failure of the peripheral choices within 5 s after nose withdrawal from the central port was also treated as miss error, though it occurred rarely (<5%). There was no punishment in any error trials and the next trial was allowed to be initiated after ITI. In the no-signal trials, animals need to wait for 0.2–0.6 s without stimulus and another 0.5–1 s to get a reward from the central port. There was no cue to distinguish the initial delay (0.2–0.6 s) and reward delay (0.5–1 s). Thus, animals did not have any external clue to differentiate the signal trials from the no-signal trials, except for the presentation of the signal itself. In the FC trials, the shutter was closed 0.5 s after stimulus presentation onset, and the rats were forced to choose either the left or right port (Fig.1a right). In cases where no stimuli were presented in FC trials, the animals were never rewarded.

Session B was the same protocol with session A described above except that only single stimulus difficulty was used. We applied the stimulus contrast level of 70-85% accuracy in FC trials in session A.

### Surgery

Rats were anesthetized with 2.5% isoflurane before surgery, and it was maintained throughout surgical procedures. We monitored body movements and hind leg reflex and adjusted the depth of the anesthesia as needed. An eye ointment was used to keep the eyes moistened throughout the surgery. Subcutaneous scalp injection of a lidocaine 1% solution provided local anesthesia before the incision. A craniotomy was performed over the anterior part of the right V1 (AP - 6.36 to −7.32 mm, ML 3.2 mm relative to the bregma, 0.2 to 0.4 mm below the brain surface) and right PPC (AP −3.8 mm, ML 2.5 mm relative to the bregma, 0.2 to 0.4 mm below the brain surface) and a custom-designed electrode was vertically implanted using a stereotactic manipulator. A stainless-steel screw was placed over the cerebellum and served as the ground during the recordings. The mean response of all electrodes was used as a reference. During a week of postsurgical recovery, we gradually lowered the tetrodes to detect unit activities in the V1 and PPC. Electrode placement was estimated based on the depth and was histologically confirmed at the end of the experiments.

### Histology

Once the experiments were completed, the rats were deeply anesthetized with sodium pentobarbital and then transcardially perfused with phosphate-buffered saline and 4% paraformaldehyde. The brains were removed and post-fixed in 4% paraformaldehyde, and 100 *μ* m coronal sections of the brains were prepared to confirm the recording sites.

### Electrophysiological recordings

A custom-designed electrode composed of two eight-tetrodes (tungsten wire, 12.5 μm, California Fine Wire, Grover Beach, CA) was used for the simultaneous recordings of V1 and PPC. The tetrodes were individually covered by a polyimide tube (A-M Systems, Sequim, WA), were placed at a 100 μm separation, and typically had an impedance of 120–1000 kΩ at 1 kHz. The signals were recorded with Open Ephys (Cambridge, MA) at a sampling rate of 30 kHz and bandpass filtered between 0.3 and 6 kHz. The tetrodes were lowered approximately 40μm after each recording session.

### Spike sorting and screening criteria of units

All analyses were performed using MATLAB (MathWorks, Natick, MA). To detect single-neuron responses, the spikes were manually clustered with MClust (A.D. Redish) for MATLAB. Only neurons met the following criteria were included for further analyses: (1) units with sufficient isolation quality (isolation distance ≧ 15); (2) units with reliable refractory periods (violations were less than 1% of all spikes); and (3) units with sufficient mean firing rates in the 1 s after the cue onset (> 1 Hz).

### Behavioral data analysis

Spatial choice accuracy was the percentage of the correct peripheral port choices in trials where either peripheral port was chosen upon the presentation of the peripheral stimulus (Fig.1b). The miss rate was the percentage of central port choices in trials where visual stimuli were presented in 3C trials (Fig.1c). The correct rejection rate was the percentage of central choices in trials where visual stimuli were not emitted in 3C trials (Fig.1d). Reaction time was defined as the duration from stimulus presentation onset to nose withdrawal from the central hole. Trials with the reaction time of less than 100 ms were considered invalid and were excluded from the calculation of the spatial choice accuracy as they were too early to have been responses to the stimulus^42^. All error bar data are presented as mean ± SEM. In all violin plots, it combined a boxplot with the kernel density estimation procedure. The boxplot inside the violin showed the quartile, whisker, and median values as white dots (Extended Data Fig.2a-h).

We classified the following three choice types based on the subjects’ detection performance (Fig.1f): (1) Hit was the trials where subjects performed peripheral choice in 3C or peripheral choice before shutter closure in FC. (2) Miss+ was the trials where they nose-poked in central port over 0.5s and they could choose the correct peripheral choice in forced-choice by shutter closure. (3) Miss− was error choice in a forced choice. Trials with miss responses in 3C were excluded from the analysis of comparison across choice types because we cannot label the trials into Miss+ or Miss-.

To estimate the impact of task parameters on behavioral performance, we conducted a generalized linear model (GLM) analysis for spatial choice (Left/Right) and hit/miss choice (Fig.1g-h). Because behavioral performance in 3C trials for the spatial choice was near perfect (90% choice accuracy), we used the identity function as the link function. In other models, we used the logit function as the link function. For spatial choice GLM analysis, we prepared trials in which they performed peripheral choices. Task parameters included with binary predictors of the stimulus (1 was stimulus presence, and 0 otherwise), peripheral previous reward (1 was rewarded, 0 otherwise), and central previous reward (1 was rewarded, 0 otherwise). The model was fit with these predictors. For hit/miss choice GLM analysis, we first prepared trials in which stimulus is present. Task parameters included with binary predictors of peripheral previous reward (1 was rewarded, 0 otherwise), central previous reward (1 was rewarded, 0 otherwise), and previous failure (1 was failed, 0 otherwise). Then we fitted the model to behavioral performance as the same procedure as spatial choice GLM analysis.

### Time-locked kernel regression and visual sensitivity

To identify visual responsive neurons, we used a time-locked kernel regression approach (Fig.1i, j, Extended Data Fig.4a-c). In this approach, the firing rate of recorded neurons is described as a linear sum of task predictors aligned to task events. In this study, we considered the stimulus onset and reaction timing kernels. According to this kernel, the predicted firing rate *f_n_*(*t*) for a neuron *n* is described as

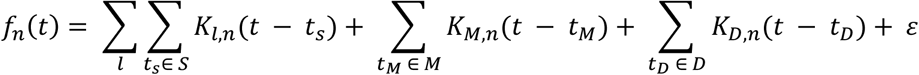

in which *l* represents the stimulus direction (ipsi or contra), *S, M, D* represents the set of times to cover each predictors window, and *K_l,n_, K_M,n_, K_D,n_* represents the stimulus, pre-motor, and pre-choice kernels for neuron *n*. The stimulus kernels cover the window 0 – 0.3s from stimulus onset and the pre-motor and choice kernels cover the window −0.3 – 0 from reaction timing (central withdrawn timing). The pre-motor and the choice kernel are identical except that the latter is designed to be sensitive to the choice directions. To fit each firing rate to the model, the firing rate was binned into 0.01-s bins and then smoothed with a causal Gaussian filter with a standard deviation 0.03s.

To determine whether each neuron is sensitive to visual response, we prepared a predictor matrix with full kernels (real design matrix) and matrix of which target kernel is shuffled within whole-time points (shuffled design matrix, Extended Data Fig.4a). We then fit the model with each design matrix to recorded firing rates and calculated the *F*-statistic of the nested model comparison test in which the reduced model was the model without that target kernel included. This procedure using a shuffled design matrix was repeated 1000 times for statistical measurement. If the *F*-statistic of the real design matrix scored high value compared to the 95% percentile of the shuffled design matrix, the neuron was deemed selective for the target kernel (Extended Data Fig.4c). In the case of neurons that were selective to the contra-stimulus kernel, we labeled stimulus preferring neurons and the other neurons were labeled stimulus non-preferring neurons.

### Selective responses to previous outcome and choice type

Neurons were classified as preferring to the previous outcome and choice type based on their mean response during the pre-stimulus epoch (−0.2 – 0s window from stimulus onset) (Extended Data Fig.5). The modulation index was computed using a receiver operating characteristic (ROC) analysis, which is to provide a measure of the difference of neural firing distribution between trial types^43^. We compared distribution between the trials in previous rewarded and unrewarded or in Hit and Miss+. The previous outcome and choice type selectivity was obtained from the area under the ROC curve (AUC) and defined as 2 × (AUC–0.5) ranging from –1 to 1. In our analysis, a positive value indicated a neuron selectively fired to the previous outcome or Hit trials, and a negative value indicated to suppress (Extended Data Fig.5a, b). A value of zero indicated the absence of selective responses. To determine statistical significance (*P* < 0.05), we used permutation tests (2,000 iteration).

### Spike train analysis

We recorded 966 neurons (V1: 515, PPC: 436 neurons) from 62 sessions in seven rats. Unless otherwise stated, the activity of each neuron was binned at 0.01s, smoothed with a causal Gaussian filter with a standard deviation 0.03s to obtain the temporal profile of each neural activity.

For visualization (Fig.1l-n) and analysis, firing rates were z-scored relative to trial-by-trial baseline rates (from the window −0.4 to 0s).

### Statistics

We evaluated the statistical significance in the analysis using data resampling with a bootstrapping procedure^51^. We estimated the *P* value for the bootstrapping procedure by computing the ratio (1+*X*) / (*N*+1), where the number *X* indicates overlapping data points between the two distributions, and the number *N* indicates iterations. Since we used 1,000 bootstraps, two distributions with no overlap resulted in *P* < 0.001, and two distributions with *x%* overlap resulted in *P* ≈ *x* /100.

### State space analysis

For the State space analysis, we used neurons with at least 20 available trials for each Hit and Miss+ condition. To characterize the population structure and the temporal pattern among all neurons during the analysis window (−0.1 – 0.15s from stimulus onset), z-scored firing rates were formatted as 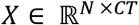, where *N* is the total number of neurons, *C* is the total number of conditions (choice types), and *T* is the number of analyzed time points. Principal component analysis (PCA) was used to reduce the dimensionality of the population from the number of neurons to three principal components (PCs). Each PC represents a weighted combination of individual neural activity, which summarized population activity.

To estimate the difference of each neural trajectories at each time points across choice types in the PC space, we prepared the dataset by bootstrapping 1000 times with different subsets of ten trials for each choice type and shuffled dataset which shuffled the weight of each PC for each neuron. We then calculated the sensitivity index (*d’*) for each PC as follows.

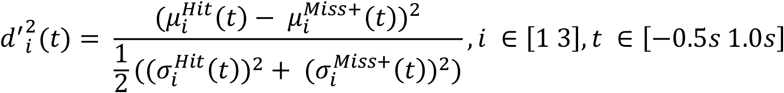

where 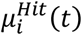 and 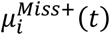 are the mean value of the *i*-th PC at time *t* in Hit and Miss+ trials, respectively, and 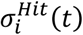 and 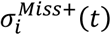 are the standard deviation of the *i*-th PC at time *t* in Hit and Miss+ trials, respectively. We used the sensitivity in the first three PCs subspace defined as the square root of *d’*^2^(*t*) (Fig.2d, g, i), as follows.

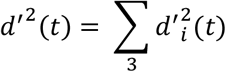

### Classification (decoding) analysis

For classifiers, we used the support vector machines (SVM) with a linear kernel function implemented using the LIVSVM library (version 3.24)^52^. All population classification was analyzed on the concatenated neural activity of individual neurons. Because the number of simultaneously-recoded neurons was low in our dataset, we constructed ‘pseudo-trials’ by randomly extracting trials from desired conditions for each neuron^53^. Data were divided into non-overlapped training/validation and test data (30% trials). To avoid overfitting, models were trained exclusively on the train/validation data set. Hyperparameter such as *C* regularization weight was determined by optimization to minimize loss of 10-fold cross-validation by grid search manner (searched range 2^−9^ – 2^15^).

For two-class classification such as classification of the stimulus (presence/absence) (Extended Data Fig.4d) and choice type (Hit/Miss+) (Fig.3a-d, h-i), we used firing rate in stimulus window (0 – 0.2s from stimulus onset), and stimulus and pre-stimulus window (−0.2 – 0s from stimulus onset) for each individual neuron, respectively. We then concatenated neural activity as described above and performed training and predictions.

For classification by adding-in subsets of neurons (Fig.3b, d), the whole population was divided into 20 equal subpopulations in the order of their individual classification performance, and then the classifier (SVM model) was built by the same procedure described above in each subpopulation.

For classification of the simultaneously-recoded population (Fig.4a-b), we first extracted sessions with at least five neurons in each region and at least 20 trials in each condition (20 sessions). We trained the classifier as the same procedure described above and predicted the test data. In the decorrelated population in V1 and PPC (Fig.4b), the trials were shuffled within trials in each neuron. We then calculated the classification accuracy of real data and the decorrelated population as the same procedure described above.

To measure the contributions of each neuron for the choice types (Fig.3g), we calculated weights of classifiers in each classifier. We compared distributions of weights for different neural types, i.e., stimulus and previous outcome selective neurons. For statistical significance, we performed a one-way ANOVA test with Tukey-Kramer post hoc comparisons.

### Population coding index

To assess how much information on a population activity improved by the information of individual neurons, we first calculated the difference between the classification accuracy of population activity and the highest individual neuron classification accuracy in that population in each population size (Fig.3e, classification accuracy in population activity – individual max accuracy). We then simply estimated the sum of the difference in all population size as population information against individual information (i.e., population coding index) (Fig.3f).

### Stability (time-resolved classification analysis)

To estimate the stability of population coding, we applied a time-resolved classification analysis where the classifiers are trained and tested with unique time samples (Fig.4h-i). Each classifier trained at time *t_trained_* can also be tested on its classification ability to predict choice type at time *t_tested_*. For visualization, we scored <0.5 classification accuracy as 0.5, and ranged 0.5 – 1.0 classification accuracy.

To estimate the stability of population coding, we calculated Pearson’s correlation coefficients between neuronal weights of each classifier at time *t_trained_* and *t_tested_* (Fig.4c). To quantify time-resolved decay of population activity pattern, we used Pearson’s correlation coefficients in 0 – 0.2s from time *t_trained_* (Fig.4e). For comparison between populations, we used t-test for each time point in V1 and PPC (Fig.4f, top), and one-way ANOVA followed by post-hoc Tukey tests for subpopulation (Fig.4f, below).

### Noise correlation

The noise correlation was defined as the correlation coefficients between the noise of neuron pairs to a given visual stimulus using z-scored firing rate for a given visual stimulus. We applied this analysis to spike counts from a 200ms window during −0.4 – 0.4s from stimulus onset (Fig.4c-d, f, Extended Data Fig.8). In this analysis, we selected neurons with at least twenty trials for each choice type (Hit and Miss+). For the statistical significance of noise correlation between choice types, we performed a paired t-test for each time epoch (Fig.4d, f, Extended Data Fig.8).

### Cell-type classification

To classify the putative fast-spiking (FS) interneuron and regular-spiking (RS) excitatory neuron, we calculated trough to late peak, and firing rate for each recorded unit (Extended Data Fig.7). Next, we determined cell-types by clustering the units in the dimension of the parameters using the k-means algorithm (k = 2). After clustering units, we defined cluster which is less trough to late peak compared to other cluster as putative FS interneurons and the other as RS excitatory neurons.

## Extended data

**Extended Data Fig 1.**
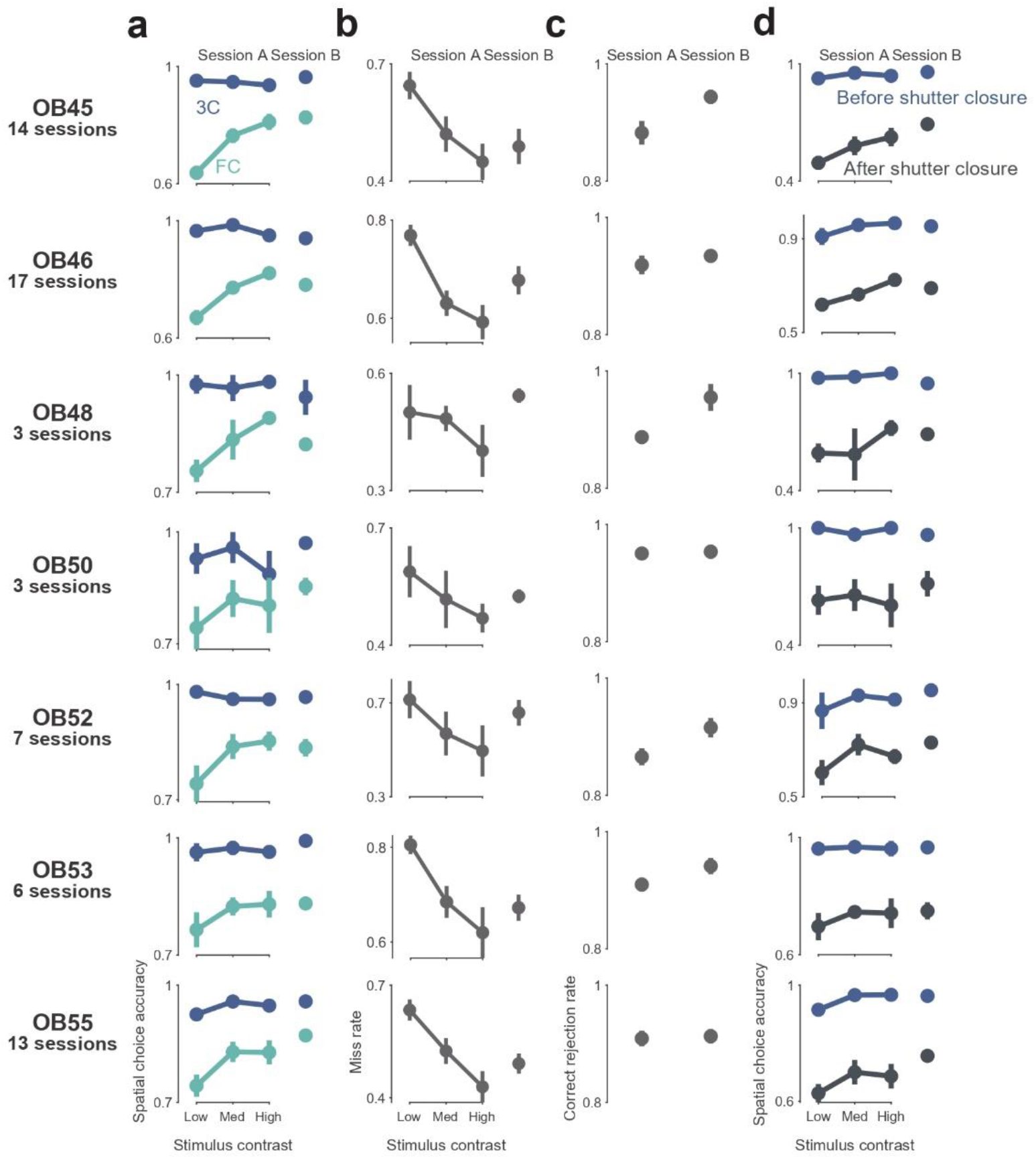
Behavioral performance in visual detection task for each subject. **a**, Spatial choice accuracy in 3-choice (3C) and forced-choice (FC) trials for each subject. The vertical axis indicates accuracy and the horizontal axis indicates three graded stimulus contrast in session A and one contrast in session B. Each subject showed above 90% accuracy in 3C trials and graded accuracy with stimulus contrast in FC trials **b**, Miss rate in 3C trials for each subject in session A and B. Each subject showed a decrease of miss rate with an increase of stimulus contrast. **c**, Correct rejection rate for each subject in session A and B. Each subject showed about 90% accuracy in both sessions A and B. **d**, Spatial choice accuracy in FC trials for each subject in session A and B. Colors correspond to choice timing (blue: before shutter closure, black: after shutter closure).

**Extended Data Fig 2.**
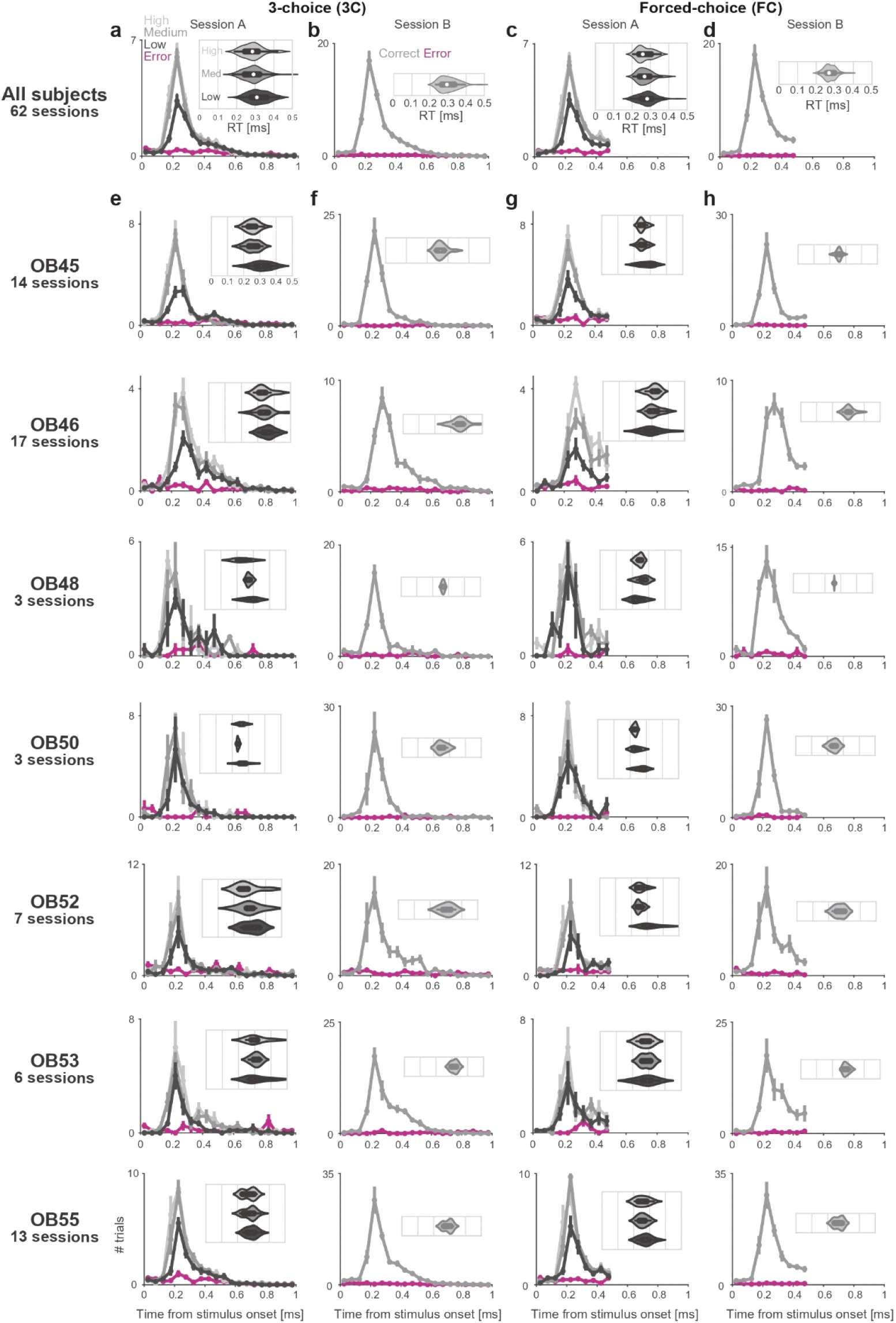
Reaction time in each subject. **a,** Distribution of reaction timing in 3C trials of all subjects in session A. Colors correspond to visual contrast and error. Inset shows violin plots of reaction timing in each stimulus contrast. White circles indicate the median value and the end of the thick line indicates quartiles. **b,** Same as in **a**, but in session B. **c,** Same as in **a**, but in FC trials. Because of shutter closure, reaction timing was not defined after 0.5s from stimulus onset. **d,** Same as in **b**, but in session B. **e-h,** Same as in **a-d** for each subject.

**Extended Data Fig 3.**
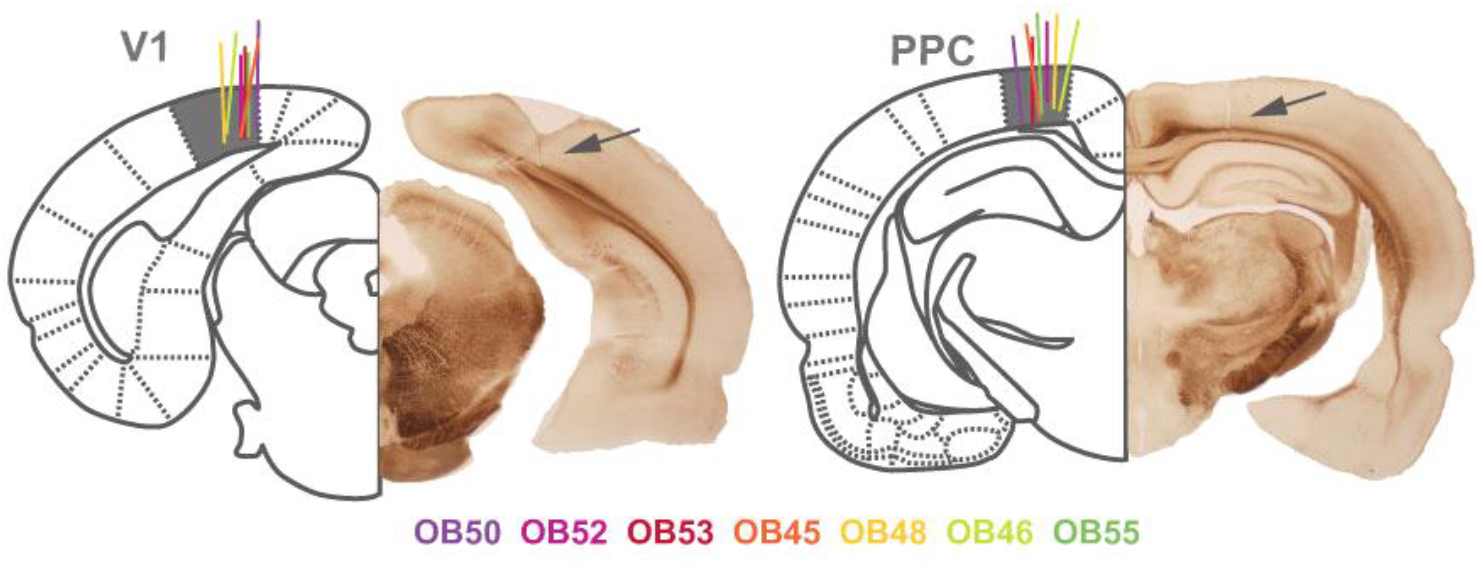
Recoding sites in V1 and PPC. Coronal section indicating recoding sites in V1 and PPC (arrow). We recorded neural activity in V1 and PPC simultaneously from seven rats. Each color corresponds to the subject ID.

**Extended Data Fig 4.**
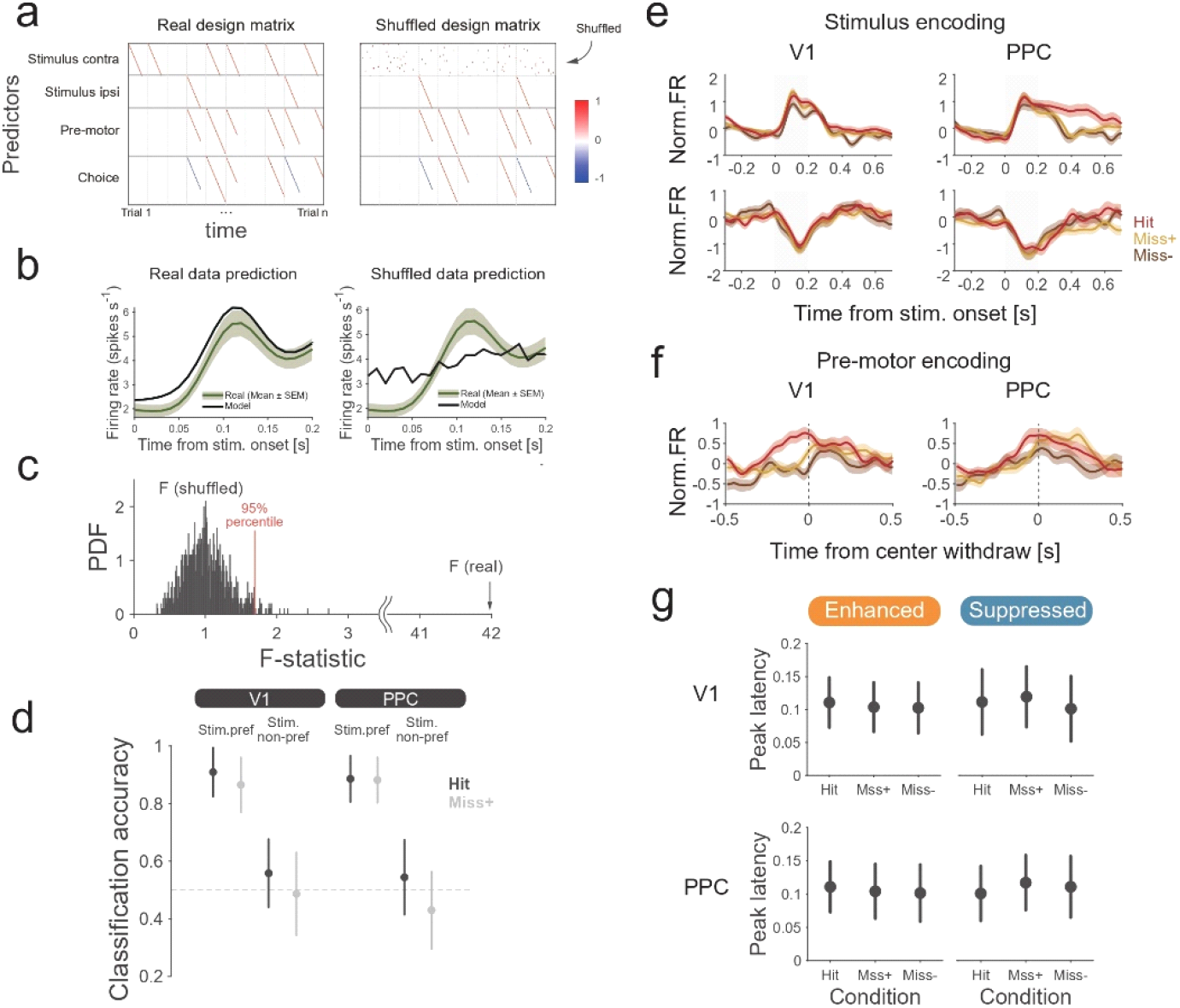
Time-locked kernel regression. **a,** Structure of predictors (kernels) design matrices. Rows of these matrices represent task variables (predictors) which take −1, 0, or 1 value for time points relative to the appropriate time offset from the specific task events. In the shuffled design matrix, the values of the target predictor (for example, contra-stimulus in the Extended Data Fig.a) were shuffled through the rows (time). **b,** Example fit of individual neurons by using real design matrix (left) and shuffled design matrix which target kernel is contra-stimulus (right). Black lines show fitted data by each model. The firing rate of the example neuron is shown as mean ± SEM (Green line and shaded area). **c,** Distributions of F-statistics calculated from the shuffled model (1000 data points). If the F-statistics calculated from the real model shows high value compared to the 95% percentile of the distributions, the neuron is sensitive to the target kernel. **d,** Classification accuracy of the stimulus (presence/absence) in Hit and Miss+ trials in stimulus preferring and non-preferring population was shown as mean ± standard deviation (STD). **e,** Trial-averaged firing rates of the stimulus encoding (preferring) neurons defined by time-locked kernel regression (target kernel is contra-stimulus) in V1 and PPC (same as in Fig.1l-m) aligned to stimulus onset. **f,** Trial-averaged firing rates of the pre-movement encoding neurons in V1 and PPC (same as in Fig.1l-m) aligned to reaction timing. **g,** Peak-latency of stimulus preferring neurons in Enhanced and Suppressed types in each choice type were shown as mean ± SEM.

**Extended Data Fig 5.**
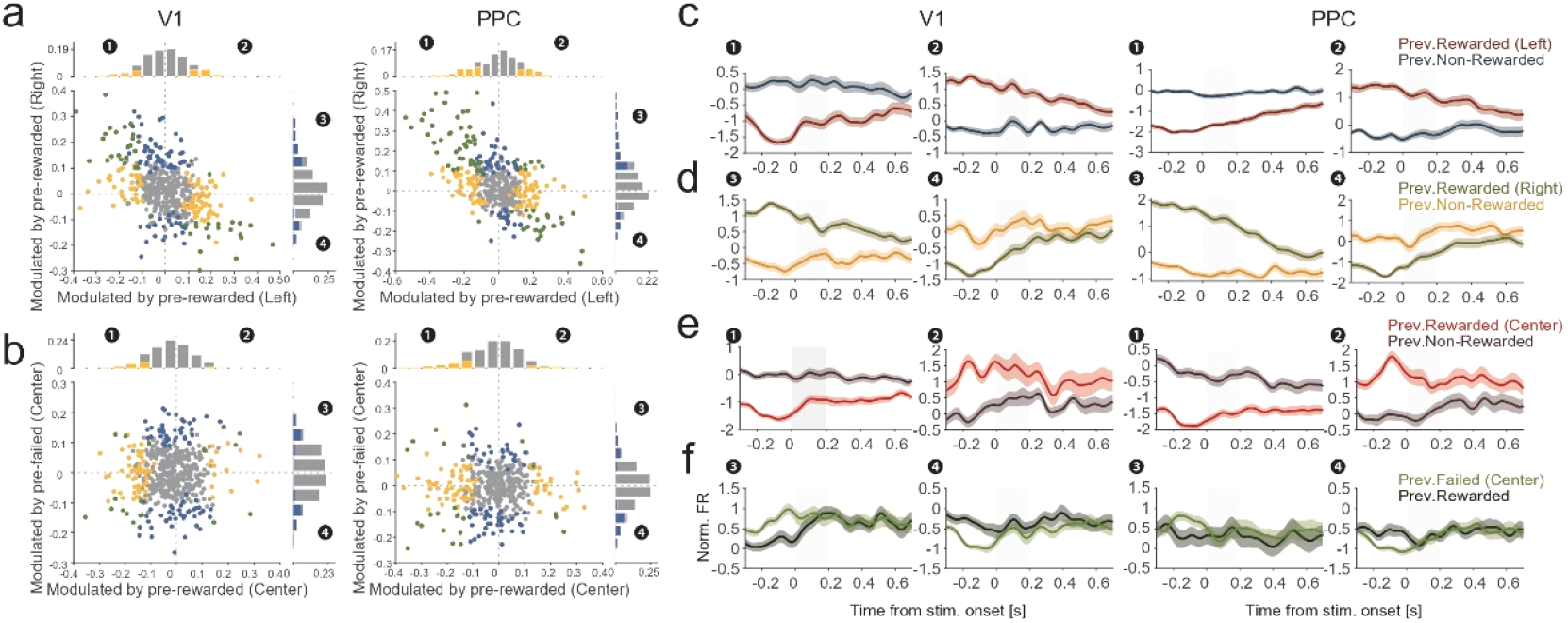
ROC analysis for selectivity of previous outcome. **a**, Scatter plot and histograms of the previous outcome modulation in V1 and PPC. Colored bars and scatters indicate the neurons with statistical significance for the previous outcome modulation (p < 0.05, permutation test). **c-f**, Time-averaged z-scored firing rates of neurons that are selective to the previous outcome aligned to stimulus onset. The gray shaded area indicates the stimulus presentation timing. Error bars indicate SEM across neurons.

**Extended Data Fig 6.**
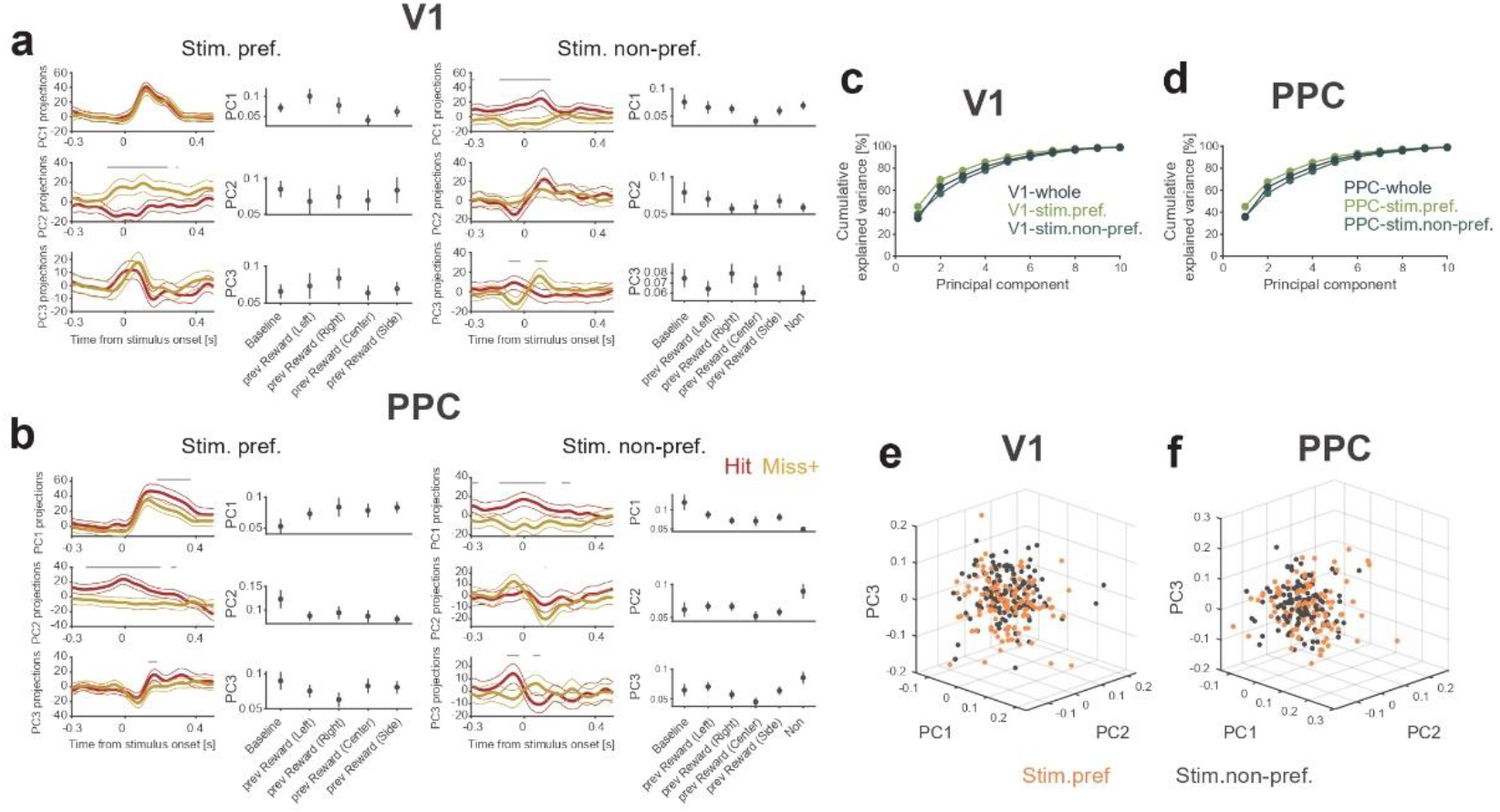
State space analysis in stimulus preferring and non-preferring population. **a-b**, PC projections for each choice type (left) and neural weights of each PC (right) in stimulus preferring and non-preferring population in V1 (**a**) and PPC (**b**). Principal components are computed on condition-averaged responses. **c-d**, The cumulative variance explained by the first 10 PCs calculated from whole, stimulus preferring, and non-preferring population. The variance is calculated over choice types and times. **e-f**, Distributions of weights by first three PCs. The weights of stimulus preferring and non-preferring neurons were distributed in the PC weight space in V1 and PPC.

**Extended Data Fig 7.**
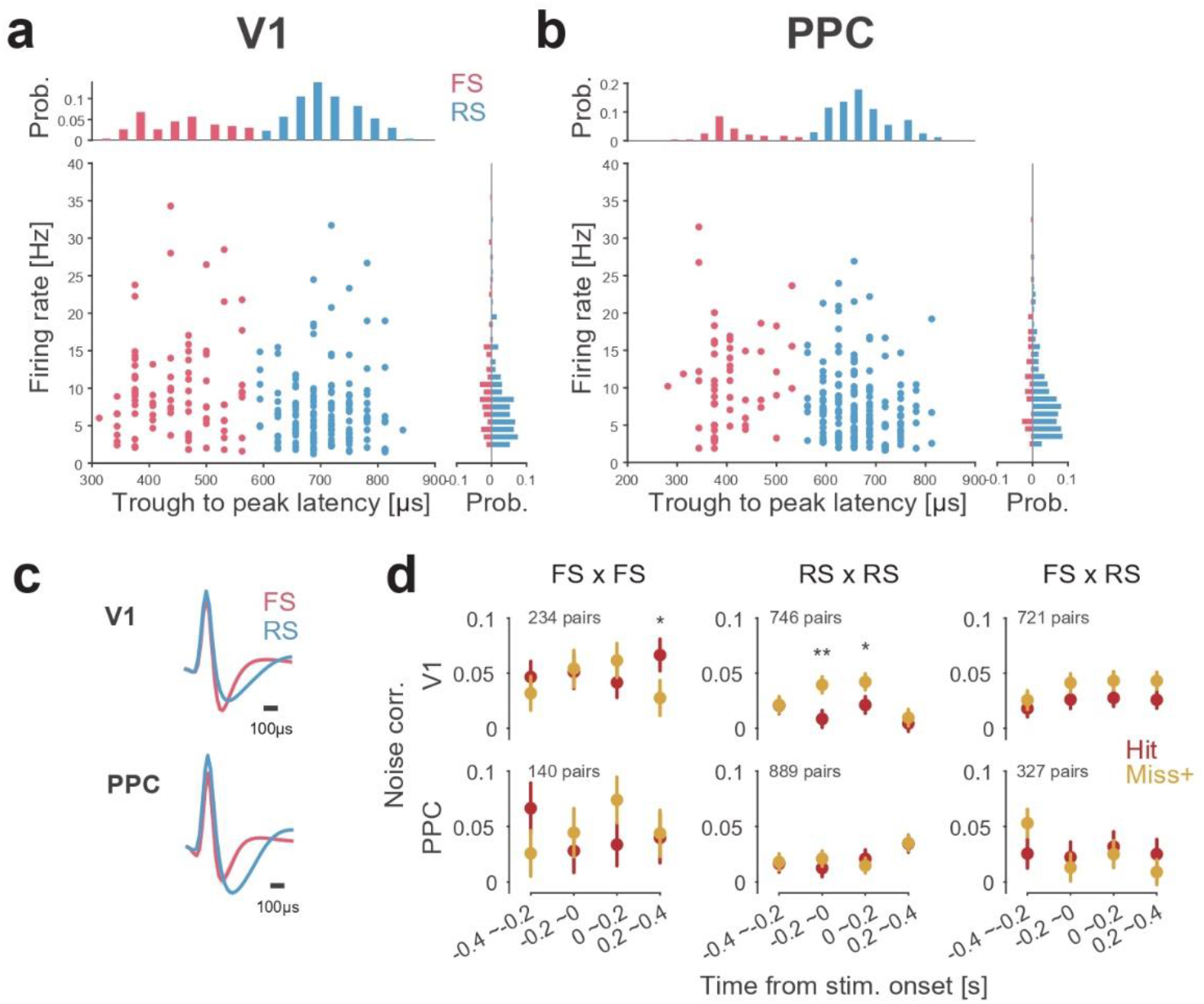
Cell-type classification. **a-b,** Scatter plot and histograms of the trough to (late) peak latency and firing rates in V1 and PPC. Each color corresponds to putative fast-spiking (FS) interneurons (pink) and regular-spiking (RS) neurons (blue). **c,** Averaged waveforms of putative FS and RS neurons in V1 and PPC. **d**, Mean noise correlation in each cell type pairs. *p<0.05, **p<0.01, t-test.

